# A Birth-Death-Migration Model for Alignment-Free Phylogeny Estimation using *k*-mer Frequencies

**DOI:** 10.1101/2023.12.05.570306

**Authors:** Siam Habib, A.T.M Mizanur Rahman, Md. Mohaiminul Islam, Khandaker Mushfiqur Rahman, Atif Rahman

**Affiliations:** Department of Computer Science and Engineering, Bangladesh University of Engineering and Technology, Dhaka

**Author notes:** These authors are co-first authors (Siam Habib, A.T.M Mizanur Rahman).

**Keywords:** phylogeny estimation, alignment-free, *k*-mer frequency, likelihood, birth-death-migration process

## Abstract

Estimating phylogenetic trees from molecular data often involves first performing a multiple sequence alignment of the sequences and then identifying the tree that maximizes likelihood computed under a model of nucleotide substitution. However, sequence alignment is computationally challenging for long sequences, especially in the presence of genomic rearrangements. To address this, methods for constructing phylogenetic trees without aligning the sequences i.e. alignment-free methods have been proposed. They are generally fast and can be used to construct phylogenetic trees of a large number of species but they primarily estimate phylogenies by computing pairwise distances and are not based on statistical models of molecular evolution. In this paper, we introduce a model for *k*-mer frequency change based on a birth-death-migration process which can be used to estimate maximum likelihood phylogenies from frequencies of *k*-mers in genomic sequences of species in an alignment-free approach. Experiments on real and simulated data demonstrate the efficacy of the model for likelihood based alignment-free phylogeny construction.

## Introduction

The phylogeny estimation problem is concerned with constructing an evolutionary tree of a set of species given their phylogenetic data and is considered one of the grand challenges in biology. In the past, physiological and morphological traits were used to determine phylogenies but in modern times we primarily rely on genomic sequences. From a computational perspective, we can model the genomic sequences as strings, and thus the phylogeny problem takes as input a set of species and strings associated with each species, and returns a tree that “best” fits the data (genomic sequences). We decide which tree is the best tree by imposing an objective function on the tree space (the set of phylogenetic trees) and we wish to find the tree that maximizes the objective.

There are two common approaches to phylogeny estimation based on how objective functions are constructed, distance based (Cavalli-Sforza and Edwards, 1967; Fitch and Margoliash, 1967; Kidd and Sgaramella-Zonta, 1971) and character based (Edwards, 1963; Fitch, 1971). In distance based approaches, a distance metric is defined over the set of all available values for phylogenetic data. The goal is to place species with small distances between their phylogenetic data close together in the tree and species with large distances farther apart. This is done by computing a distance matrix containing the pairwise distance between the phylogenetic data for every pair of species and then constructing a weighted tree that minimizes the “deviation” between the distance matrix and the distances between each pair of species in the constructed tree.

In character based approaches, instead of constructing a distance matrix from the phylogenetic data, a “character matrix” is constructed. The individual characters are represented using the columns of the matrix and the rows represent the species. We assume that the characters change according to some rule. We try to place two species with similar characteristics close to one another in the phylogenetic tree. This is done by defining an objective function on the set of all phylogenetic trees given the character matrix and trying to find a tree that optimizes it. Two such candidates for objective functions are the parsimony function (Edwards and Cavalli, 1964; Fitch, 1971) and the likelihood function (Edwards and Cavalli, 1964; Fisher, 1921, 1922). Among all the methods of phylogenetic reconstruction, likelihood based ones have been observed to be the most successful (Felsenstein, 1978, 1981; Kuhner and Felsenstein, 1994; Yang, 1994).

In likelihood based approaches, a statistical model is defined over how characters transition over time. Different positions of characters are usually assumed to be independent of one another and how a character changes with time is modeled as a memoryless stochastic process i.e. a continuous time Markov process (Ross, 2014). Given the character matrix, a statistical model and a rooted weighted tree, one can thus find the likelihood of that tree (given the character matrix). The goal in this case is to find a tree with the maximum likelihood.

The models used in likelihood based phylogeny estimation are based on statistical observations about molecular evolution, i.e. how genomic data change over time. Some of the notable ones among such models are Jukes-Cantor69 (Jukes et al., 1969), Kimura80 (Kimura et al., 1980), Kimura81 (Kimura, 1981), Felsenstein81 (Felsenstein, 1981), HKY85 (Hasegawa et al., 1985), and GTR (Tavaré, 1986). To use any such model in practice, first a multiple sequence alignment must be constructed from the genomic data and then the multiple sequence alignment is used as the character matrix for the model. Computation of multiple sequence alignment is a hard problem in computer science both in theory (Wang and Jiang, 1994) and in practice (Kemena and Notredame, 2009; Thompson et al., 1999, 2011).

To overcome the computational cost of aligning the sequences, there has been a rising interest in alignment-free methods, where phylogenetic trees are constructed without explicitly aligning sequences. A multitude of alignment-free methods for phylogeny construction have been developed recently and there is considerable variation in their approaches (Haubold, 2014). Some notable alignment-free methods include kSNP (Gardner and Hall, 2013; Gardner and Slezak, 2010; Gardner et al., 2015), co-phylog (Yi and Jin, 2013), mash (Ondov et al., 2016), andi (Haubold et al., 2015), multi-SpaM (Dencker et al., 2020).

Alignment-free methods also have benefits other than avoiding the computational cost of multiple sequence alignments. Sequence alignment generally assumes that changes to the genomic sequences only happen due to small local changes. But, in reality, alterations to genomic sequences also occur due to large scale genomic rearrangements such as reversals, translocations, duplications, deletions. Alignment based phylogeny estimation methods may lead to incorrect results in the presence of such rearrangements which can potentially be remedied by alignment-free methods.

While alignment-free methods seem very promising due to their scalability, performance, and ability to take into account non-local changes, most of them are distcance based whereas the best performing phylogeny construction tools in the alignment-based paradigm are still likelihood based. Although, Höhl and Ragan (Höhl and Ragan, 2007), and Zahin et al. (Zahin et al., 2019) presented Bayesian and likelihood based alignment-free methods of phylogeny estimation using the absence or presence of *k*-mers, they rely on existing models for binary traits for likelihood computation, and do not explicitly model the process of changes to *k*-mer frequencies.

In this paper, we introduce a linear birth-death model with constant positive migration to model how the *k*-mer frequency profile of a species changes with time, paving the way for a method for constructing phylogenetic trees that tries to get the best of both worlds i.e. that is both alignment-free and likelihood based. Unlike (Höhl and Ragan, 2007) and (Zahin et al., 2019), our model takes into account the frequencies of different *k*-mers adding further distinction between *k*-mers that are present. Our method counts the frequencies of different *k*-mers in the genomic data of each species and exploits the fact that if the *k*-mer frequency profile of two species are similar they must be close to each other in the evolutionary tree. The model allows us to compare how likely a tree is, given the *k*-mer profiles of each species and thus combines the advantages of alignment-free methods and the mathematical soundness of likelihood based methods.

## Materials and Mehods

### Birth-Death-Migration Process

A birth-death-migration process (Bailey, 1991) is a continuous time Markov process where there are infinite states, each state is labeled with a non-negative integer. Every state *i* has a transition to state *i*+1 with transition rate *λ*_*i*_ that models population growth (a birth or a positive migration) and every state *i>* 0 has a transition to state *i*−1 with transition rate *µ*_*i*_ that models a population decrease (death or negative migration). Figure 1 shows the Markov chain for a birth-death-migration process.

**FIG. 1.**
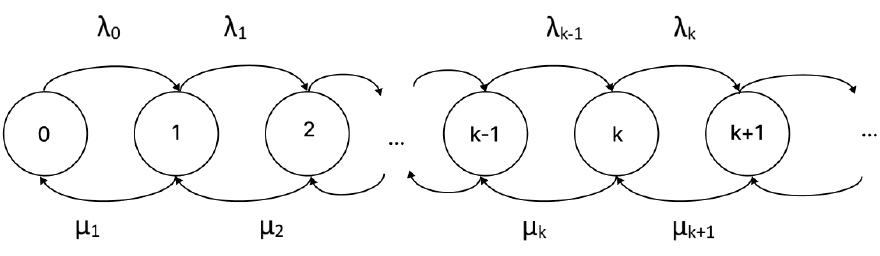
A birth-death-migration process

Note that in a birth-death-migration process, a population growth event can happen in two ways, through a birth or through a positive or incoming migration. We will denote the birth rate at state *i* with *b*_*i*_ and the positive migration rate at state *i* with 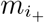. Then, we can write

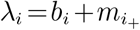

Similarly in a birth-death-migration process, a population decrease can happen in two ways, through a death or through a negative or outgoing migration event. We will denote the death rate at state *i* with *d*_*i*_ and the negative migration rate at state *i* with 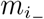. So, we can write

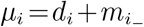

When modeling populations in many scenarios, the rate of birth, the rate of death, and the rate of negative migration can be assumed to be linear in the size of the current population i.e.,

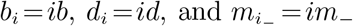

where *b, d* and *m*_−_ are the rates of birth, death and negative migration respectively when the population size is 1.

### Modeling evolution of *k*-mer frequencies

Consider a set of *k*-mers and their occurrences in the genomic sequence of a species. We can consider occurrences of different *k*-mers as distinct populations. Then, sizes of the populations represent the *k*-mer frequencies. The changes in *k*-mer frequencies because of small scale substitutions, insertion-deletions, and genomic rearrangements parallel changes in the sizes of the populations due to births, deaths, and migration. We can model how the population changes with time with a birth-death-migration process. Births model increase of a *k*-mer frequency due to a duplication event. Deaths model the decrease of a *k*-mer frequency because of a deletion event. Migrations are the increase or decrease of a *k*-mer frequency due to substitutions of a single or a few nucleotides, small insertion-deletions, or at the breakpoints of genomic rearrangements.

For example, Figure 2 illustrates the process considering three populations, the occurrences of CGTG, CTTG, and CTTA. Initially, the sequence was ATCGTGCGTGTACTTG and CGTG, CTTG, and CTTA had populations of sizes 2, 1, and 0, respectively. First, a deletion event happens that deletes an occurrence of CGTG and reduces the population size to 1 (in the second frame from the top). This reduction in population can be modeled as a death event.

**FIG. 2.**
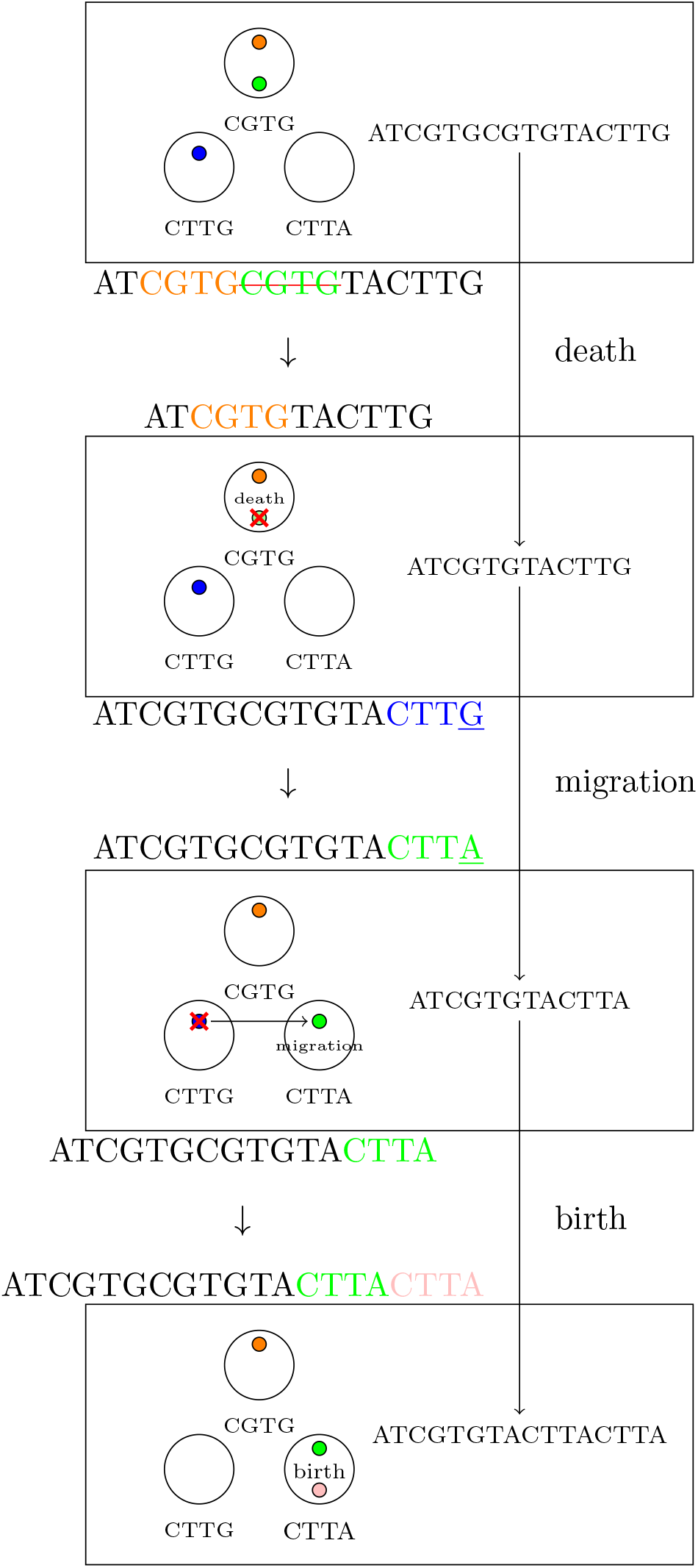
Change of *k*-mer frequencies with time

Then, a substitution of a single nucleotide happens that changes an occurrence of CTTG to CTTA. This can be thought of as a negative migration in the population of CTTG and a positive migration for CTTA (in Frame 3). Finally, in Frame 4, a duplication event happens for the occurrence of CTTA that increases the population size to 2. This is modeled as a birth event. Thus, we can model the following biological phenomena with birth-death and migration as follows:

#### Duplication

In a duplication event (Frame 4 of Figure 2), occurrences of some *k*-mers get duplicated thus increasing the population sizes of the *k*-mers. We can model this as a birth. Assuming that duplication is a memoryless stochastic process that is equally likely to happen at all sites in the genome, the rate of birth for any *k*-mer population at any given time is proportional to the size of the population i.e. birth rate is linear.

#### Local Alterations

We refer to substitutions of a single or a few nucleotides, small insertions-deletions, and changes at the breakpoints of genome rearrangements as local alterations (Frame 3). When a local alteration happens, occurrences of a few *k*-mers change to others. This has two effects. It acts as negative migration for one set of populations and positive migration for others. By the same argument for gene duplication, the rate of negative migration is linear. But positive migration is not. The rate at which positive migration to a *k*-mer *l* happens is proportional to the number occurrences of *k*-mers that are similar to *l* but not *l*. If we make the simplifying assumption that the rate of transition from any *k*-mer *p* to another *k*-mer *q* due to local alteration is roughly equal, we can model positive migration by a constant (during stationarity).

#### Deletion

In a deletion event, an occurrence of some *k*-mers get deleted (Frame 2). By similar arguments to the ones we presented above, this can be modeled with a linear death rate.

Thus we can model how the *k*-mer frequency of a single *k*-mer changes with time using a birth-death-migration process with linear rates of birth, death and negative migration, and a constant rate of positive migration i.e.

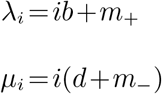

The model is simplified by denoting *µ* = *d*+*m*_−_ and by using the symbol *m* and *λ* in place of *m*_+_ and *b*, giving us the following equations:

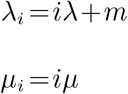

We make further assumptions that the rates *λ, µ* and *m* are the same for all *k*-mers (for a fixed value of *k*) and independent of what the *k*-mer is i.e. mutations and rearrangements occur with the same rate at all sites in the genome. Furthermore, we assume that the *k*-mer frequency for different *k*-mers are independent and identically distributed.

### Overview of alignment-free likelihood based phylogeny estimation

The components in a conventional alignment based phylogeny estimation scheme using likelihood is shown in Figure 3a whereas Figure 3b illustrates the proposed alignment-free likelihood based method. The components and the modifications are described below:

**FIG. 3.**
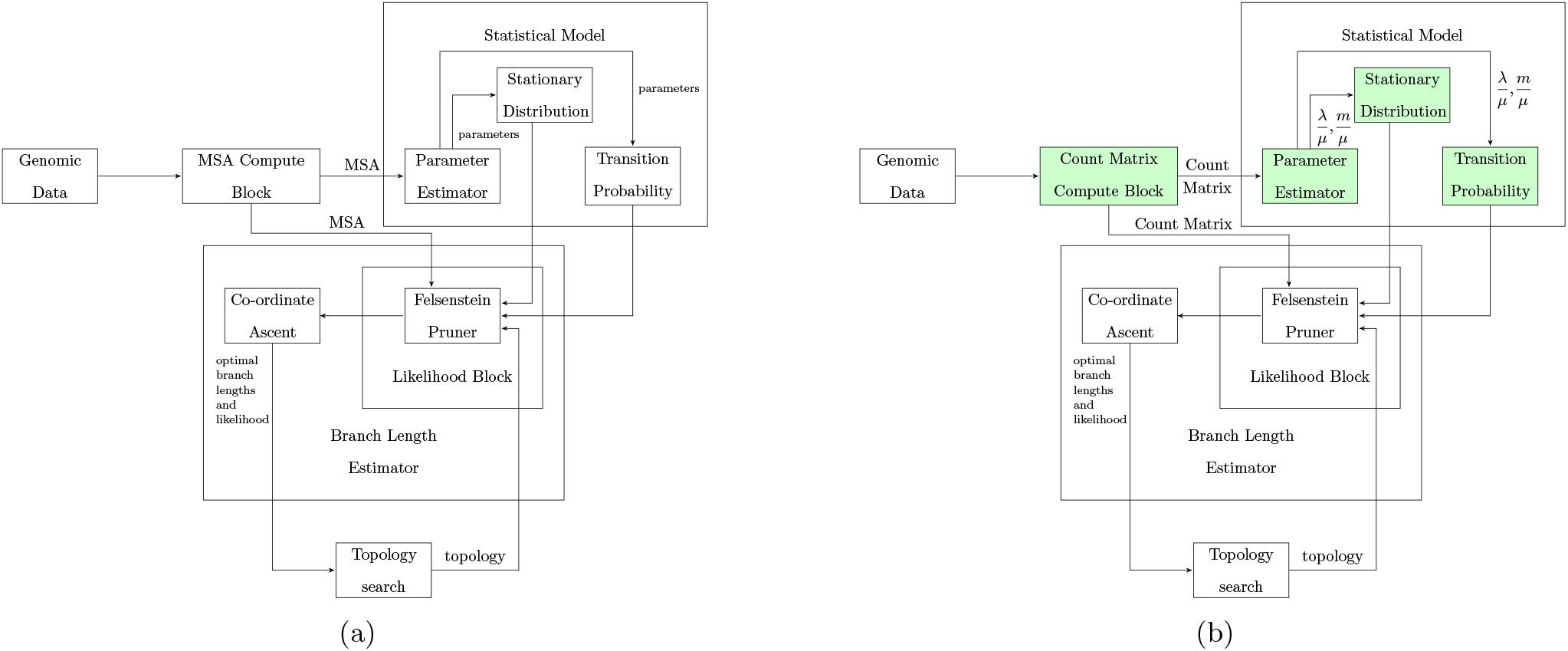
Overviews of a) a conventional alignment-based and b) the proposed alignment-free likelihood based phylogeny estimation methods. The modules that distinguishes the two approaches are highlighted in green.

#### Character matrix computation

First, the character matrix computation module takes as input the genomic data and computes a character matrix. For conventional likelihood based methods, this is done using multiple sequence alignment. In our approach, it computes a count matrix where each column represents counts of a *k*-mer in each of the species.

#### Parameter estimation

The parameter estimator block takes as input the character matrix and estimates the parameters of the parametric statistical model the likelihood based phylogeny estimator is based on. In our case, it estimates the parameters for the birth-death-migration process.

#### Likelihood computation

A likelihood computation block takes as input the statistical model, the parameters of that model, the character matrix, and a weighted tree, and computes the likelihood of that tree given the parameters and the character matrix. This block uses Felsenstein’s Pruning algorithm (Felsenstein, 1973, 1981) to compute the likelihood in linear time (in the size of the tree). In our alignment-free setting, the algorithm uses the stationary distribution and the transition probabilities of our birth-death-migration model.

#### Branch length estimation

The branch length estimator block takes as input a tree topology, the statistical model, and the parameters of that model as input and finds tree branch lengths that maximize the likelihood for that tree. This module uses numerical optimizers and repeatedly calls Felsenstein’s Pruning algorithm to estimate the branch lengths.

#### Tree topology search

The tree topology search unit searches over the tree topology space using some kind of local search algorithm. This module picks a tree topology and calls the branch length estimator to maximize the likelihood of that topology and then explores neighboring topologies. Our model can be incorporated in existing tree search methods to find the maximum likelihood tree.

We now describe how we compute the expressions in detail in the following sections.

### Count Matrix Computation

The count matrix computation module takes as input the genomic sequences and computes a count matrix by counting the number of occurrences of different *k*-mers and their reverse complements in the given sequences. We have used Jellyfish (Marcais and Kingsford, 2011) to implement this module.

Before generating the count matrix, we must choose the value of *k* (the length of *k*-mers) and decide which *k*-mers to include in the count matrix. The reason we do not include all *k*-mers of length *k* is two folds. First, allowing all *k*-mers for a fixed size might make the matrix too large (4^*k*^*/*2 *k*-mers for length *k*). Second, it is assumed in our model that the *k*-mer frequencies for any two different *k*-mers are independent and identically distributed. This will not be the case if many *k*-mers that are close to one another are present in the matrix. So, to reduce the dependency, we uniformly sample *n*_*k*_ *k*-mers from all *k*-mers of length *k* (assuming that a *k*-mer and its reverse complement is equivalent).

Thus our count matrix computation module has two hyper-parameters: (i) *k*, the length of the *k*-mers that we will count, and (ii) *n*_*k*_, the number of *k*-mers to sample. This is also the number of columns in the count matrix. We select the value of the hyper-parameters by picking the pair (*k*,*n*_*k*_) that produces the highest entropy (Shannon, 1948; Sherwin, 2010) *k*-mer existence matrix, a 0-1 matrix where the entry of the *i*th row and *j*th column is 1 if and only if the *j*th *k*-mer exists in the genome sequence of the *i*th species. The entropy of the *k*-mer existence matrix is defined as the average of the entropy of each column in the matrix and the entropy of column *j* is defined as 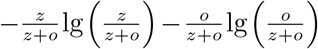 where *z* and *o* denote the numbers of zeros and ones in the *j*th column of the *k*-mer existence matrix respectively.

We choose (*k,n*_*k*_) by looping over the integer interval [5,17] for *k* and setting 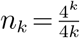. As there are 3*k* +1 strings that are at most one edit distance apart from any *k*-mer, if we let *n*_*k*_ to be greater than 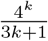 we are guaranteed to have two closely related *k*-mers in our count matrix. We therefore round up 3*k* +1 to 4*k* to set 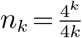. We also restrict *n*_*k*_ to be within 5000 and 50000 and clip *n*_*k*_ appropritely if it is not. This is done to make sure that *n*_*k*_ is not too small or too large.

### Computing Stationary Distribution

We can find the expression for the stationary distribution of our linear birth-death-migration model by using the balance equations:

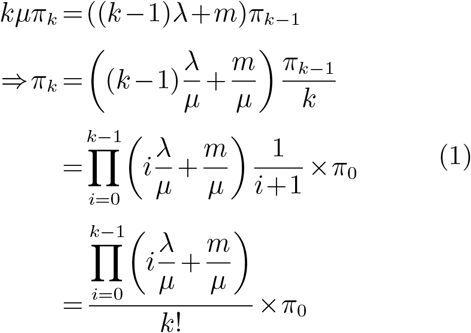

By using 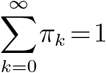 and Equation 1, we get:

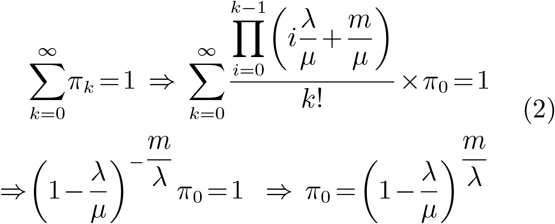

By substituting the value of *π*_0_ in Equation 1, we find the expression for *π*_*k*_ for all *k* ≥ 0.

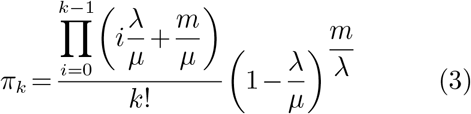

Thus we can find the value of *π*_*k*_ for any *k* in 𝒪 (*k*) operations. Note that from Equation 3, we can see that for the stationary distribution to exist, *λ* ≤ *µ*. In practice, we are actually more interested in computing the log value of the stationary probabilities instead of the stationary probabilities themselves.

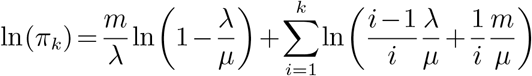

### Parameter Estimation

First note that, the parameters of our model *λ, µ*, and *m* are dependent on the unit of time and thus any linear scaling of the terms will yield equal likelihood for the same tree (where the branch lengths are scaled down by the same factor). Thus instead of using *λ, µ* and *m* as parameters, we can use 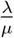 and 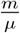. We will denote our parameters 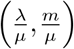 using *θ* from now on.

Now assume that we are given a *k*-mer count matrix **M**_*cnt*_. From the *k*-mer count matrix we can generate a count array **c** where *c*_*i*_ is defined as the number of occurrences of the integer *i* in the entire *k*-mer frequency matrix, **M**_*cnt*_. Let *N* be the maximum entry in **M**_*cnt*_. Then the length of the count array **c** is *N* +1 and contains the indices 0 to *N*. Under the assumptions that the entries of the *k*-word count matrix were sampled independently from the stationary distribution we can find the probability of the count array **c** given the parameters 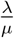 and 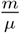 using the following:

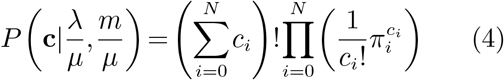

where *c*_*i*_ denotes the entry in the *i*th index in the count array **c** i.e. number of *k*-mers that have frequency *i*.

Using Equation 4, we can find the log likelihood of our parameters *θ* as follows:

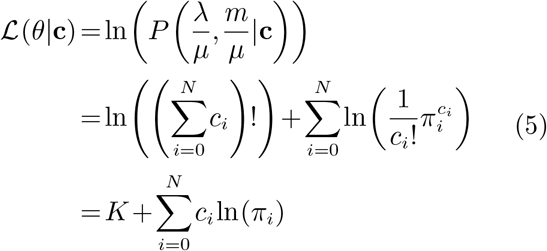

where *K* is a constant dependent on the data (the count matrix which is constant). Using Equation 5, we can find the expression for the gradient of the log-likihood:

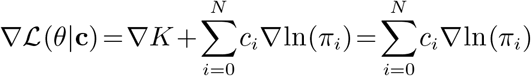

We can also find the Hessian matrix of the log-likelihood function:

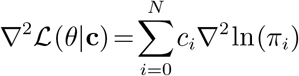

Let us further simplify our notation by using the symbols *x* and *y* respectively to denote the parameters 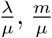. Then we can find the expressions for the gradient and the Hessian matrix for ln(*π*_*k*_) for all indices *k*.

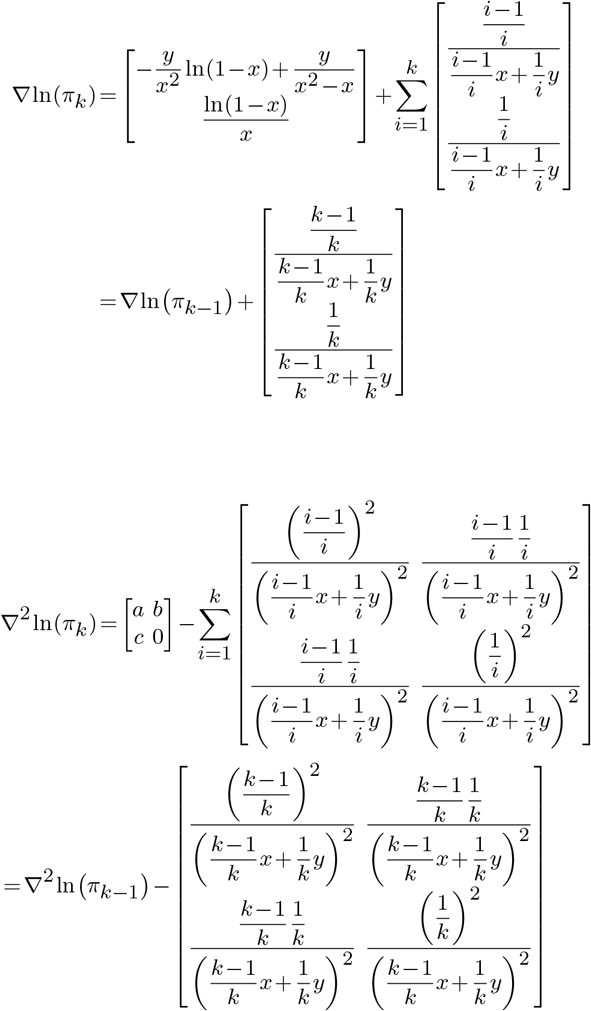

where

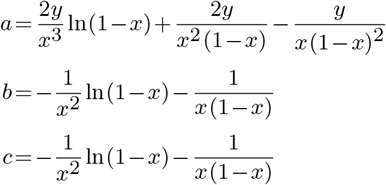

Notice that for the stationary distribution to exist, we must have that 0 *<x<* 1 and *y* ≥ 0. Thus, we have to find the most likely values for *x* and *y* under the constraints that 0 *<x<* 1 and *y* ≥ 0. The constrained region for our problem forms a rectangular region which is unbounded on one side. Finding the maximum value of a function in such a rectangle can be solved using the trust-region method (Conn et al., 2000). We used scipy’s trust-constr (scipy.org, 2023a, b) method to maximize the log likelihood of the parameters given the count array. Scipy’s trust-constr implementation is based on (Byrd et al., 1999) and (Lalee et al., 1998) who themselves implement (Byrd, 1987), (Omojokun, 1989) and (Byrd et al., 1999).

### Branch Length Estimation

Given a fixed tree topology and the model parameters, finding the optimal branch lengths that maximize the likelihood of the tree topology exactly is considered hard for most phylogenetic models. Usually, branch lengths are estimated using co-ordinate ascent (Wright, 2015) based numerical methods. Co-ordinate ascent is a numerical algorithm that maximizes an objective function *f* : ℝ^*n*^ → ℝ in an iterative manner. In each iteration, the algorithm picks a co-ordinate and optimizes that co-ordinate assuming all other co-ordinate values are fixed (Wright, 2015).

In the context of our problem, we want to maximize the likelihood of the tree given the topology *T* and parameters *θ*. Thus we may consider our likelihood function ℒ_*T,θ*_ : ℝ^|*E*|^ → ℝ to take an |*E*| dimensional vector as input, where |*E*| is the number of branches in the tree. We will denote the vector of all branch lengths as **x**. We initialize **x** with random non-negative integers. In each iteration, we loop over all the edges and update their branch lengths with the most likely length of that edge while keeping the rest of the branch lengths fixed.

Solving the problem of finding the most likely length of a branch while keeping rest of the branch lengths fixed is usually solved using numerical methods such as gradient ascent (Cauchy et al., 1847; Polyak, 1987), Newton-Raphson method, Brent’s method (Brent, 1973; Dekker, 1969), trisection search, golden section search (Kiefer, 1953). We used scipy’s `minimize scalar` function with method set to `bounded` (scipy.org, 2023c).

### Likelihood Computation

To compute the likelihood, we used Felsenstein’s Pruning algorithm. It queries the statistical model for the transition probabilities over the branch lengths. Let *S* be a continuous time Markov chain with a finite number of states. If *n* is the number of states in *S* then the rate matrix *Q* is an *n×n* matrix. We can compute the transition matrix function *P* (*t*) of *S* using *P* (*t*) = *e*^*Qt*^.

However, in the case of our birth-death-migration model, this can not be used because the birth-death-migration process has infinite states. Note that the above is true regardless of whether the CTMC is finite or not. It is just no longer possible to compute the matrix efficiently. Therefore, we approximated the transition probability function for our model by generating an approximate rate matrix *Q*^′^ by bounding the number of states to some large value. This is done by computing the stationary distribution of the *k*-mer counts using our estimated parameters and finding the largest value *i* that has a higher stationary probability than some small cut-off probability (10^−8^).

## Results

In this section we present the results of our experiments. We evaluate the accuracy of parameter estimation, and assess the effectiveness of our model in finding the actual branch lengths and tree topologies. We also use our model to analyze three datasets from AFproject (Zielezinski et al., 2019).

### Parameter Estimation

First, we evaluate the accuracy of the estimated parameters of our birth-death-migration model. As real datasets with known parameter values are not available, we use simulated data for this. We generated 60 different synthetic datasets in batches of 10 according to our model. The datasets were generated by generating random rooted trees with random branch lengths and then simulating the evolution of *k*-mer frequencies according to linear birth-death migration models with randomly generated parameters.

Since, the parameters *λ, m*, and *µ* are not uniquely identifiable from the *k*-mer count matrices, we estimated the values 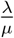 and 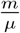, and compared with the real values. Figures 4a and 4b show scatter plots of the estimated against the true values of the parameters 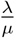 and 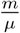, respectively. From the figures we can see that the estimated values are almost identical to the real values and the points are largely along the diagonal line. The correlation coefficient between estimated and original values are 0.9991 for 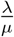 and 0.9998 for 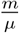. This suggests that our parameter estimation module can estimate the parameters with high accuracy.

**FIG. 4.**
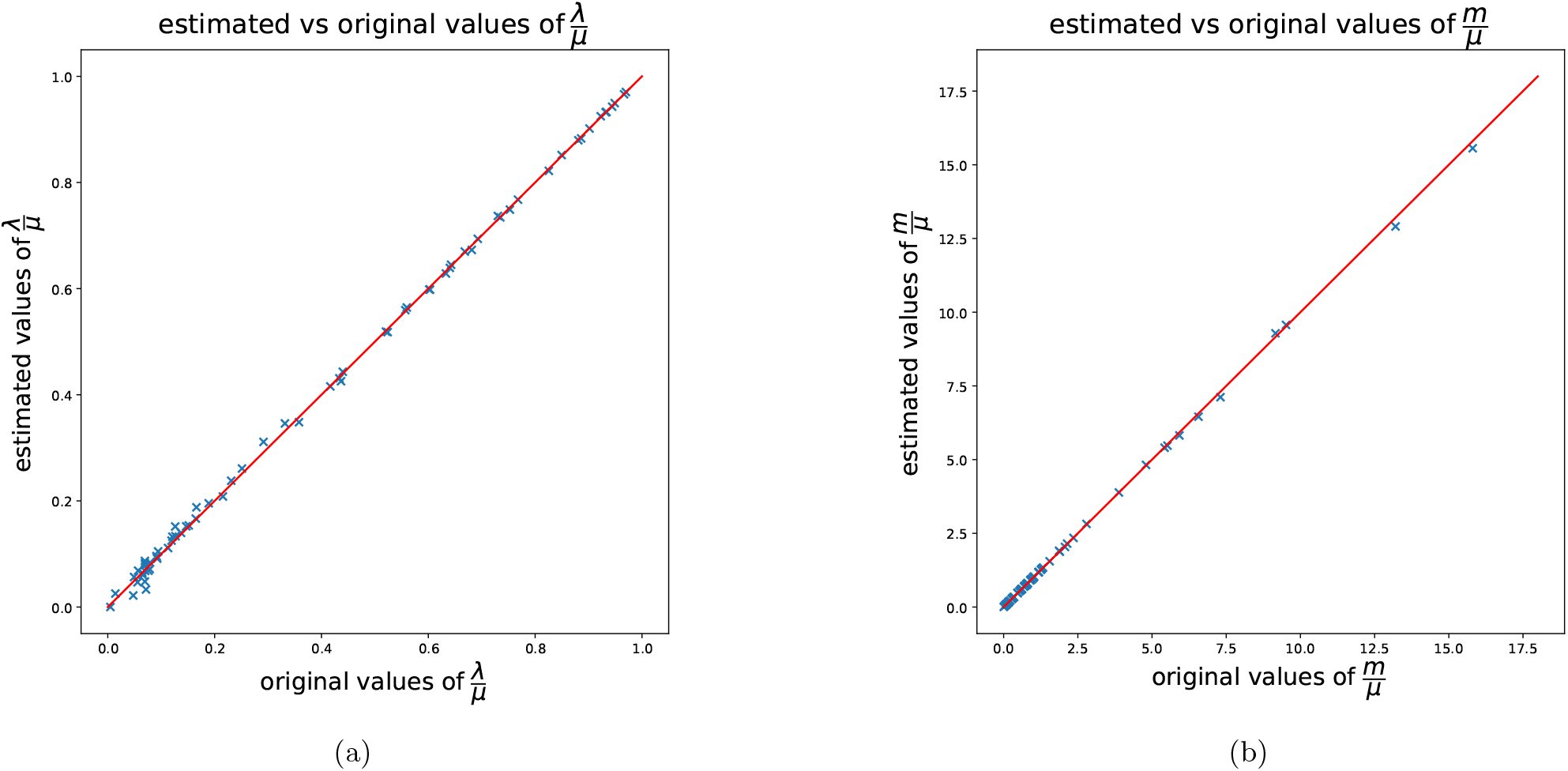
Scatter plots showing estimated parameter values vs original values for 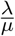 and 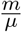.

### Branch Length Estimation

Next we assess our implementation of branch length estimation using two different batches of datasets, one simulated and one real. The synthetic datasets were generated as discussed in the previous subsection. We generated a batch of 10 trees with their birth-death-migration parameters normalized (0 *<λ<* 1 and *µ* = 1). We sampled the value of *m* uniformly from the interval (0,1). The original branch lengths of the trees are between 0.1 and 0.9 units. For the real dataset we used a seven primate dataset (Haubold, 2014). We generated the tree topology and branch lengths using the GTR+G model in RAxML-ng (Kozlov et al., 2019) and generated 7 *k*-mer count matrices using Jellyfish for all *k* in the interval [7,13].

For both datasets we evaluated our implementation of the module by providing it the original tree topology without the branch lengths. For the real dataset, we assume that the tree returned by RAxML using the GTR+G model is the true tree. We hide the original branch lengths by initializing the lengths of all the branches with a uniform random number from the interval (0,5). For the simulated datasets we provide the module with the original parameter values for the birth-death-migration process and for the real dataset, we use our parameter estimation module to estimate them before we pass it to the branch length estimation module.

Branch length distance (BLD) (Kuhner and Felsenstein, 1994) is widely used to compare estimated branch lengths with true ones. However, in our model, the branch lengths do not uniquely identify a tree even when the topology is fixed since given any tree *T*, we can find a new tree *T*^′^ with likelihood equal to that of *T* by scaling the branch lengths of *T* by a constant *K* and scaling the birth-death-migration parameters by 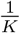 for any *K >* 0. Thus to analyze the results of our experiment we introduce the term BLDK(*T,T* ^′^,*K*) as a measure of distance between two trees.

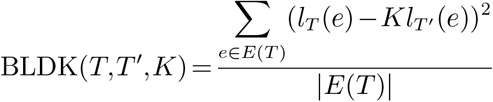

where *l*_*T*_ (*e*) and *l*_*T*′_ (*e*) denote the lengths of the branch *e* in the trees *T* and *T*^′^ respectively.

Since for the simulated dataset, we have already provided the branch length estimation module with the original birth-death-migration process parameter values, we use BLDK with *K* = 1 to analyze our results. For the real datasets, we do not know the original birth-death-migration parameters. So, we analyze the results by setting the value of *K* in such a manner that it minimizes the value of BLDK for all *K*. We can determine the value of *K* that minimizes BLDK(*T*_*real*_,*T*_*estimated*_,*K*) by taking its derivative and setting it to 0.

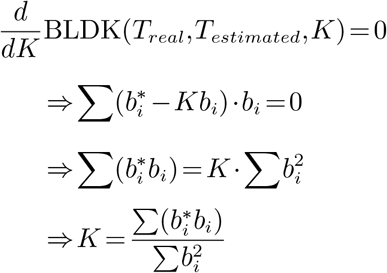

where 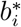 and *b*_*i*_ denote the original and estimated branch lengths of the *i*-th branch respectively.

Figure 5 shows comparisons between the BLDK of the trees estimated by our branch length computation module and the BLDK of the trees with randomly initialized branch lengths for both simulated data and real data.

**FIG. 5.**
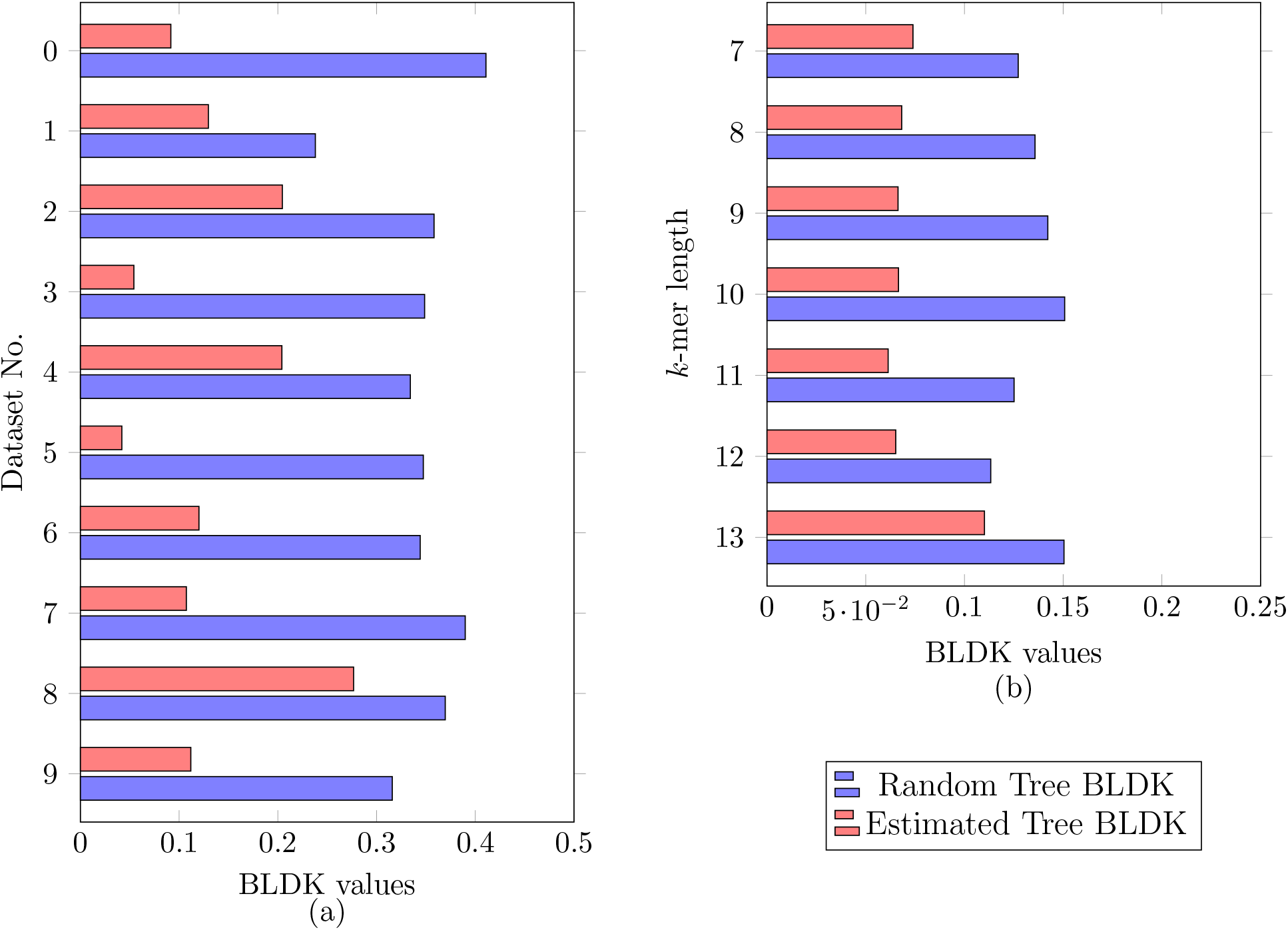
Comparison of BLDK values for trees with random branch lengths and trees with estimated branch lengths for (a) simulated data and (b) real data.

We can see that for all datasets (both real and simulated), BLDK for our estimated branch lengths are lower than those of the randomly initialized branch lengths. Additionally for the real tree, we observe the trend that as the value of *k* gets large, the value of the BLDK of the estimated tree becomes closer to the BLDK of the one initialized with random branch lengths. This is due to the fact that if the dataset is small, for large values of *k* the *k*-mer count matrix has a large number of 0s and loses most of the information. This is why we use the value of *k* that maximizes the entropy of the *k*-mer matrix. Figure 6 shows the branch lengths obtained using RAxML and our estimated branch lengths for the *k*-mer length 8 for the seven primate dataset. The choice of the value of *k* was based on maximum entropy. Since the branch lengths of both trees are not on the same scale, we re-scale the branch lengths so that the sum of the branch lengths for both trees are 1. We find that branch lengths in the two trees are similar. However, we observe that the root to leaf distance for baboon is much longer than those for other species in the tree obtained using RAxML which is anomalous (Wu et al., 2020). On the other hand, the root to leaf distances for all the species are similar to each other according to the branch lengths estimated by our method.

**FIG. 6.**
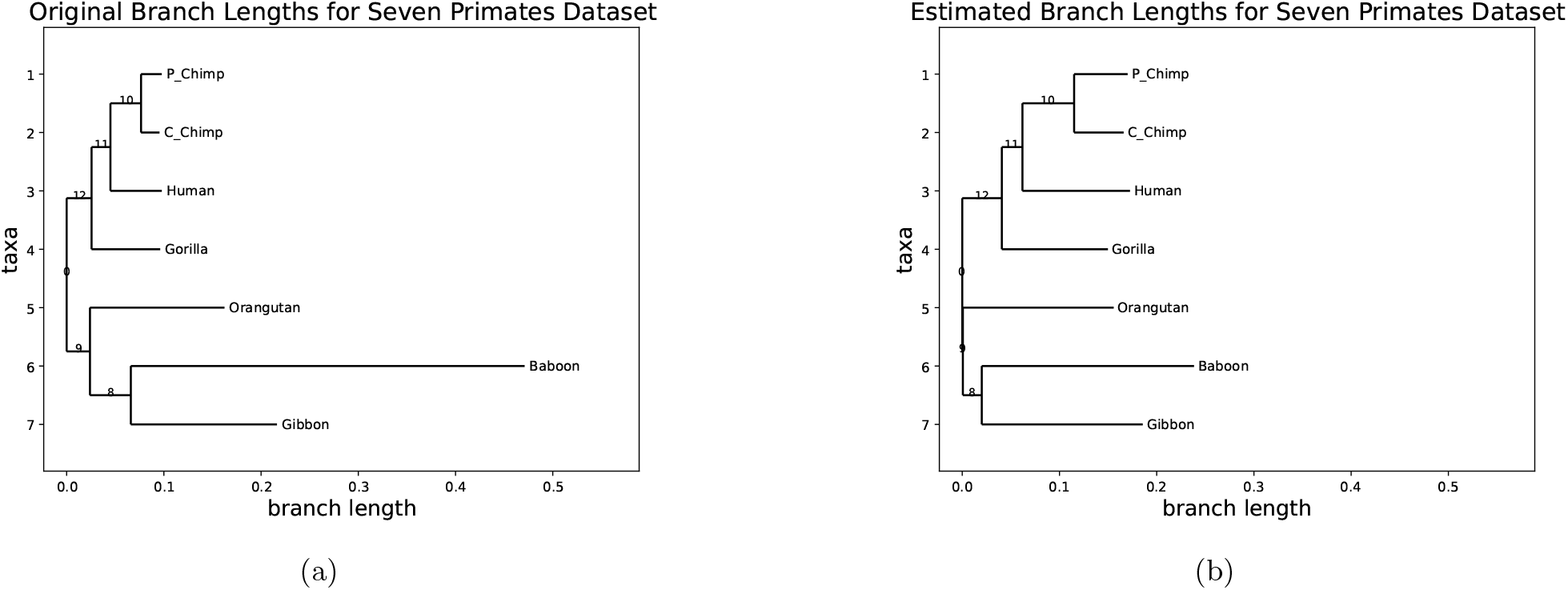
Comparison of branch lengths obtained using a) RAxML with GTR-G model and b) our birth-death-migration model for the seven primates dataset.

### Tree Topology Search

Finding the maximum likelihood tree by searching in the tree topology space requires the likelihood curve of trees to have desirable properties in the tree space. More precisely, we want the likelihood of the branch length optimized real tree to be higher than the likelihood of other branch length optimized trees. We also want trees with high likelihood to be topologically close to the real tree. However, since the tree topology space is discrete, the problem of finding an optimal tree topology is combinatorial in nature. The large number of tree topologies and the combinatorial nature of the problem makes an efficient implementation of topology search to find the optimal tree according to our model beyond the scope of this paper.

Therefore, to assess our model’s efficacy in finding the actual tree topology we perform the following experiment. Starting with the original tree, we repeatedly perturb it using the subtree pruning and regrafting (SPR) operation (Bordewich and Semple, 2005). We then compute the likelihood of the trees using our model as well as their Robinson-Foulds (RF) distances (Robinson and Foulds, 1981) from the original tree.

The experiment was done using two datasets -one synthetic and one real. The synthetic dataset was generated in a similar manner as mentioned in the previous sections. For the real dataset, we used the seven primate dataset. We assumed that the tree topology generated by RAxML for the seven primate dataset is the true tree topology.

Figure 7 shows likelihood vs RF distance for both the datasets. From the figure we can see that as the RF distance from the original topology increases, the average likelihood decreases. This demonstrates that our model assigns higher likelihood to trees topologically closer to the original tree than to the ones that are farther from it indicating its applicability to alignment-free maximum likelihood phylogeny estimation.

**FIG. 7.**
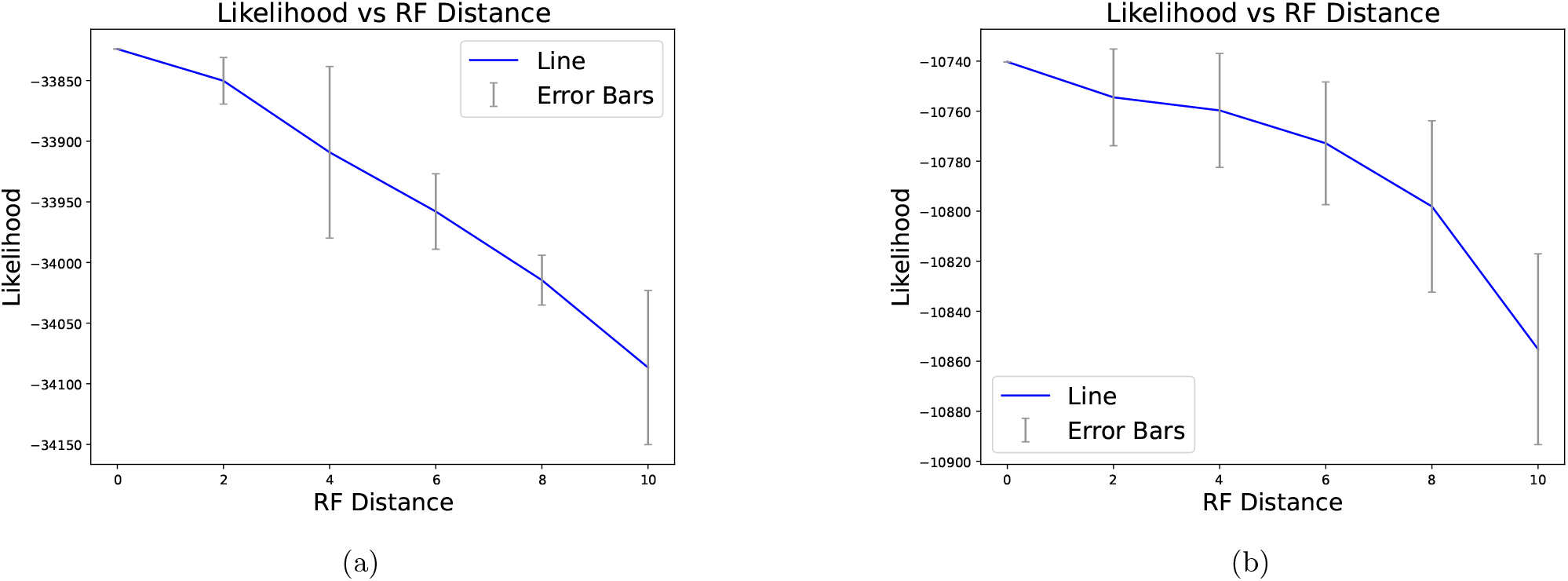
Likelihood vs RF distance of trees for a) simulated and b) seven primate datasets.

### Analysis of the AFproject Datasets

Finally, we use our model to analyze datasets from AFproject (Zielezinski et al., 2019) which has been widely used to assess various alignment-free phylogeny estimation tools. The project evaluates a multitude of alignment-free tools on a number of datasets and provides a rank-list of the tools on each dataset. AFproject also provides a “true tree” or a reference tree for each of the datasets. Here we compute the likelihood of different tree topologies already present in AFproject using our model and analyze how they compare with the ranking by AFproject.

We use three datasets for genome based phylogeny from AFproject - the *E*.*coli* dataset (Yi and Jin, 2013), the plant dataset (Hatje and Kollmar, 2012) and the fish mtDNA dataset (Fischer et al., 2013). As the number of trees available in the project for each dataset is very high, we select a subset of the trees. For each dataset, we make sure that at least one topology from ranks 1, 2 and 3 in AFproject rank-lists are selected. In addition, we select the topologies generated by some widely used tools - mash, co-phylog, skmer, kSNP3, andi, and phylonium. We include the “original” or the reference topology as well as a randomly generated topology. If any of the trees had a node with more than two children, the node was split to multiple binary nodes arbitrarily. If a tree was not rooted, a random node was selected for rooting. Models that provided their topologies in tsv or phylip format were converted to the Newick format using the Neighbor Joining algorithm (Saitou and Nei, 1987).

Tables 1, 2 and 3 show the results for the *E. coli*, the Plant and the Fish mtDNA datasets respectively. Each table shows likelihood of each topoplogy in the descending order of their likelihoods along with their RF distances from the reference tree and their rank in the AFproject ranklist. Note that the since we had to split internal nodes with more than two children into multiple children, the RF distances between trees may now differ from the value listed on AFproject.

**Table 1.** Comparison of tree topologies on the AFproject *E. coli* Dataset.

| Model | Likelihood | RF | Rank |
| --- | --- | --- | --- |
| kSNP3 | -586788.214 | 14 | 4 |
| Skmer, mash | -587693.661 | 16 | 4 |
| andi | -589409.349 | 10 | 2 |
| co-phylog | -589530.017 | 14 | 3 |
| Reference | -590045.284 | 0 | 0 |
| 8/10_DNA_TWS<br>phylonium | -590122.076 | 6 | 1 |
| FSWM/Read-SpaM | -590448.781 | 14 | 3 |
| Random | -808184.589 | 54 | — |

**Table 2.** Comparison of tree topologies on the AFproject Plant Dataset.

| Model | Likelihood | RF | Rank |
| --- | --- | --- | --- |
| Skmer | -1082511.709 | 8 | 2 |
| Reference | -1082712.868 | 0 | 0 |
| mash | -1082715.367 | 6 | 1 |
| co-phylog | -1083061.626 | 10 | 1 |
| ASW(1,11,COS) | -1083217.839 | 10 | 3 |
| kSNP3 | -1083939.777 | 16 | 5 |
| Multi-SpaM | -1084033.582 | 2 | 1 |
| phylonium | -1088458.069 | 16 | 8 |
| Random | -1121801.549 | 24 | — |
| andi | -1130283.135 | 20 | 10 |

**Table 3.** Comparison of tree topologies on the AFproject Fish mtDNA Dataset.

| Model | Likelihood | RF | Rank |
| --- | --- | --- | --- |
| ASW(1,9,EUC),<br>(4)SR(K)MER_FEM1<br>mash<br>8/10_DNA_TWS | -96915.235 | 4 | 1 |
| AFKS-kl_conditional | -96915.535 | 10 | 2 |
| AFKS-canberra | -96916.465 | 14 | 2 |
| co-phylog<br>kSNP3<br>Skmer | -96917.306 | 20 | 2 |
| AFKS-<br>k_divergence | -96917.999 | 12 | 2 |
| phylonium | -96919.240 | 12 | 3 |
| Reference | -96923.982 | 0 | 0 |
| andi | -96938.589 | 16 | 4 |
| ASW(1,3,EUC) | -98951.448 | 44 | 17 |
| Random | -101048.468 | 44 | — |

We observe that for the *E. coli* dataset, the reference topology is ranked 5th according to the likelihood computed under our model. The phylogenies constructed by kSNP3, Skmer, mash, andi and co-phylog all have likelihoods higher than that of the reference. It is worth highlighting that the reference tree for this dataset was constructed by concatenating the alignments of 2034 core genes, constructing a distance matrix and then using the Neighbor Joining algorithm (Yi and Jin, 2013; Zhou et al., 2010). The resulting tree has been subsequently used as a reference without extensive validation.

In case of the Plant dataset (Table 2), the “reference” tree got the second highest likelihood and the only topology that got higher likelihood was generated by Skmer (Sarmashghi et al., 2019). Overall for this dataset, we observe that the ranking imposed by our model is in concordance with the AFproject ranking with the exception of Skmer and Multi-SpaM.

Finally, in case of the Fish dataset, we find that although the ranking imposed by likelihood is in agreement with the ranking by AFproject, the reference tree is ranked after the trees ranked 1,2, and 3. We also observe that a large number of methods in the AFproject ranklist all generated the same topology (top most one in the AFproject ranking), and our model also placed a higher likelihood on that topology over the others.

## Discussion

In this paper, we have introduced a model for *k*-mer frequency change based on a birth-death-migration process. This model is designed to estimate maximum likelihood phylogenies from *k*-mer frequencies without aligning the sequences. Our experiments, conducted on both real and simulated data, illustrate the effectiveness of this model for alignment-free phylogenetics in a maximum likelihood paradigm. A future direction will be to explore whether some of the assumptions made in our model can be relaxed to further improve the model. In addition, analysis of the AFproject datasets reveals some discrepancies between the AFproject rankings and the likelihoods computed based on our model, warranting more investigation. Finally, the model may be incorporated in a tool that can search over the tree topology space to find a tree that maximizes the likelihood according to the model to enable alignment-free likelihood based phylogenetics.

## Code Availability

The codes underlying this article are available at https://github.com/kmerthesisgroup.

